# Aerobic H_2_ respiration enhances metabolic flexibility of methanotrophic bacteria

**DOI:** 10.1101/075549

**Authors:** Carlo R. Carere, Kiel Hards, Karen M. Houghton, Jean F. Power, Ben McDonald, Christophe Collet, Daniel J. Gapes, Richard Sparling, Gregory M. Cook, Chris Greening, Matthew B. Stott

## Abstract

Methanotrophic bacteria are important soil biofilters for the climate-active gas methane. The prevailing opinion is that these bacteria exclusively metabolise single-carbon, and in limited instances, short-chain hydrocarbons for growth. This specialist lifestyle juxtaposes metabolic flexibility, a key strategy for environmental adaptation of microorganisms. Here we show that a methanotrophic bacterium from the phylum Verrucomicrobia oxidises hydrogen gas (H_2_) during growth and persistence. *Methylacidiphilum* sp. RTK17.1 expresses a membrane-bound hydrogenase to aerobically respire molecular H_2_ at environmentally significant concentrations. While H_2_ oxidation did not support growth as the sole electron source, it significantly enhanced mixotrophic growth yields under both oxygen-replete and oxygen-limiting conditions and was sustained in non-growing cultures starved for methane. We propose that H_2_ is consumed by this bacterium for mixotrophic growth and persistence in a manner similar to other non-methanotrophic soil microorganisms. We have identified genes encoding oxygen-tolerant uptake hydrogenases in all publicly-available methanotroph genomes, suggesting that H_2_ oxidation serves a general strategy for methanotrophs to remain energised in chemically-limited environments.

## Introduction

Methane (CH_4_) is a potent greenhouse gas with a 100-year global warming potential 34 times greater than carbon dioxide (1). Aerobic methane-oxidising bacteria (methanotrophs) constitute the primary biological sink for atmospheric methane (~30 Tg annum^-1^) (2). Together with anaerobic methane-oxidising archaea (ANME), they also capture the majority of biologically- and geologically-produced methane before it enters the atmosphere (3). Relative to their global impact as greenhouse gas mitigators, methanotrophic bacteria exhibit low phylogenetic diversity and are presently limited to 22 genera in the Alphaproteobacteria and Gammaproteobacteria (4), two candidate genera (*Methylacidiphilum* and *Methylacidimicrobium*) in the phylum Verrucomicrobia (5, 6) and two representatives of candidate phylum NC10 (7, 8). Methanotrophic bacteria have been isolated from a variety of ecosystems, particularly at the oxic-anoxic interfaces where the fluxes of methane gas are high, including peat bogs, wetlands, rice paddies, forest soils and geothermal habitats (9).

Methanotrophic bacteria are classified as obligate aerobes and obligate methylotrophs. They grow by oxidising single carbon or short-chain organic compounds (10). All species oxidise methane to methanol via the soluble or particulate form of methane monooxygenase (sMMO or pMMO respectively). Methanol is then oxidised to carbon dioxide (CO_2_), yielding reducing equivalents for energisation of the respiratory chain. The proteobacterial methanotrophs generate biomass by assimilating formaldehyde, a product of MaxFI type methanol dehydrogenase activity, *via* the ribulose monophosphate or serine pathways (2). The verrucomicrobial methanotrophs, in contrast, express an XoxF type methanol dehydrogenase that oxidises methanol directly to formate (11) and generate biomass by fixing CO_2_ via the Calvin-Benson-Bassham cycle (12). To remain viable, all methanotrophic bacteria simultaneously require methane, an endogenous reductant (NAD(P)H or quinol), and an exogenous terminal electron acceptor (O2) to sustain the highly challenging methane monooxygenase reaction (CH_4_ + O2 + [NAD(P)H + H^+^]/QH_2_ → CH^3^OH + NAD(P)^+^/Q + H_2_O) and to assimilate carbon. This apparently specialist lifestyle is likely to make methanotrophic bacteria highly vulnerable to environmental changes, such as methane-limitation, hypoxia or oxidative stress. But contrary to their apparent metabolic inflexibility, it is known that methanotrophic bacteria are capable of surviving within environments where methane may be limited, variable, or restricted to atmospheric concentrations (1.7 ppmv) (13). Under these conditions low-affinity methane monooxygenase enzymes, which dominate cultivated representatives of methanotrophic bacteria (14), are incapable of methane oxidation and hence other energy sources are required for survival. Recent reports have shown that some methanotrophic bacteria can grow on simple organic acids, alcohols and short-chain alkane gases (15). While this expansion of metabolic ability supersedes the long-held paradigm that methanotrophic bacteria are obligate methylotrophs, these observations are restricted to three of the 24 described methanotrophic genera, *Methylocella, Methylocapsa* and *Methylocystis* (16). It remains unknown whether methanotrophic bacteria more commonly exploit alternative energy sources to survive periods of methane starvation.

Recent physiological and ecological studies have collectively shown hydrogen is a widely utilised energy source for microbial growth and survival across a growing range of taxa and soil ecosystems (17-19). The discovery that the dominant soil phyla including Actinobacteria (20) and Acidobacteria (18) can switch from growing on heterotrophic substrates to persisting on atmospheric hydrogen has provided a new understanding of how microorganisms survive nutrient-limited conditions. Given its ubiquity and diffusivity, hydrogen gas is an ideal energy source to support the growth or non-replicative persistence of soil bacteria. Our recent survey of hydrogenase distribution (17), metalloenzymes that catalyse the reversible oxidation of hydrogen, noted that genes encoding hydrogenases were extremely widespread. This result led us to investigate the distribution of hydrogenases in methanotrophic bacteria, and we subsequently identified genes encoding these enzymes in all 31 publicly available genomes (Figure S1). This prevalence of hydrogenase genes was surprising given the apparent specialist lifestyle of methanotrophic bacteria. Reports of hydrogenase activity in these microorganisms are scarce. Formate-dependent hydrogen production, under anoxic conditions, has been shown in cultures of *Methylomicrobium album* BG8 and *Methylosinus trichosporium* OB3b (21). *Methylococcus capsulatus* (Bath) is known to express both cytosolic and membrane-bound hydrogenases, though their physiological function has not been resolved (22). The oxidation of hydrogen in methanotrophic bacteria has been predicted to contribute reducing energy for methane oxidation (22), to recycle endogenous hydrogen produced during nitrogen-fixation (23), and to drive the non-productive oxidation of chlorinated solvents (24). However, no studies have confirmed whether hydrogen metabolism contributes to the growth and persistence of these microorganisms.

To investigate the physiological role of hydrogenases in methanotrophic bacteria, we undertook a geochemical, molecular and cultivation-based survey of a geothermal soil profile at Rotokawa, New Zealand with known methanotrophic activity (25). In this work, we isolated a thermoacidophilic methanotroph from the genus *Methylacidiphilum* and demonstrate that it oxidises hydrogen gas during both growth and persistence. Hydrogen oxidation occurred through an aerobic respiratory process mediated by a membrane-bound [NiFe]-hydrogenase. Hydrogen oxidation enhanced mixotrophic growth yields during methanotrophic growth, under both oxygen-replete and oxygen-limiting conditions, and was sustained in the absence of methane oxidation. We propose aerobic hydrogen respiration serves as a dependable mechanism for this bacterium – and potentially methanotrophic bacteria in general – to remain energised in otherwise physically-challenging and energetically-variable environments.

## Results

**Verrucomicrobial methanotrophs are abundant in environments with high levels of soil methane and hydrogen oxidation.** A molecular survey of total microbial taxa revealed bacterial and archaeal populations were consistent with acidic soil ecosystems. Methanotrophic verrucomicrobial taxa (family Methylacidiphilaceae) were the dominant OTUs identified at 10 cm depth, accounting for 47 % of all bacterial 16S rRNA gene sequences retrieved (Figure 1a). *Methylacidiphilum* spp. have been previously isolated in acidic geothermal soils in Kamchatka (26), Italy (27, 28) and New Zealand (29) (pH optima of < 3.0), and along with *Methylacidimicrobium* spp. (6), are the only acidophilic methanotrophic bacteria described. At depths between 20-50 cm, Proteobacteria, Actinobacteria and Acidobacteria were present at greater abundance than Verrucomicrobia, none of which include known acidophilic methanotrophs or hydrogenotrophs. At all depths, total microbial taxa were dominated by archaeal OTUs, classified into the order Thermoplasmatales, which is consistent with observations made previously of other acidic geothermal environments (25, 30, 31). Methane and hydrogen soil gas concentrations in this soil embankment were greatest at a depth of 50 cm and steeply decreased below detectable limits by 10 cm depth, suggesting high levels of methane and hydrogen gas oxidation was occurring in shallow depth soils (Figure 1). We consistently observed rapid oxidation of hydrogen and methane in these surface soils when incubated *in vitro* at both mesophilic and thermophilic temperatures (Figure S2). Anticipating soil gas oxidation would primarily be driven by verrucomicrobial methanotrophs, we designed PCR primers targeting the methane monooxygenase and hydrogenase genes encoded in the *Methylacidiphilum infernorum* V4 genome (32) (Table S1). qPCR analysis on soil DNA extracts confirmed the genetic capacity of Methylacidiphilaceae to oxidise methane and hydrogen. The abundance of these genes was greatest in the top 20 cm of soil, which corresponded to the zones of the lowest methane and hydrogen soil gas concentrations (Figures 1b & 1c).

**Figure 1:**
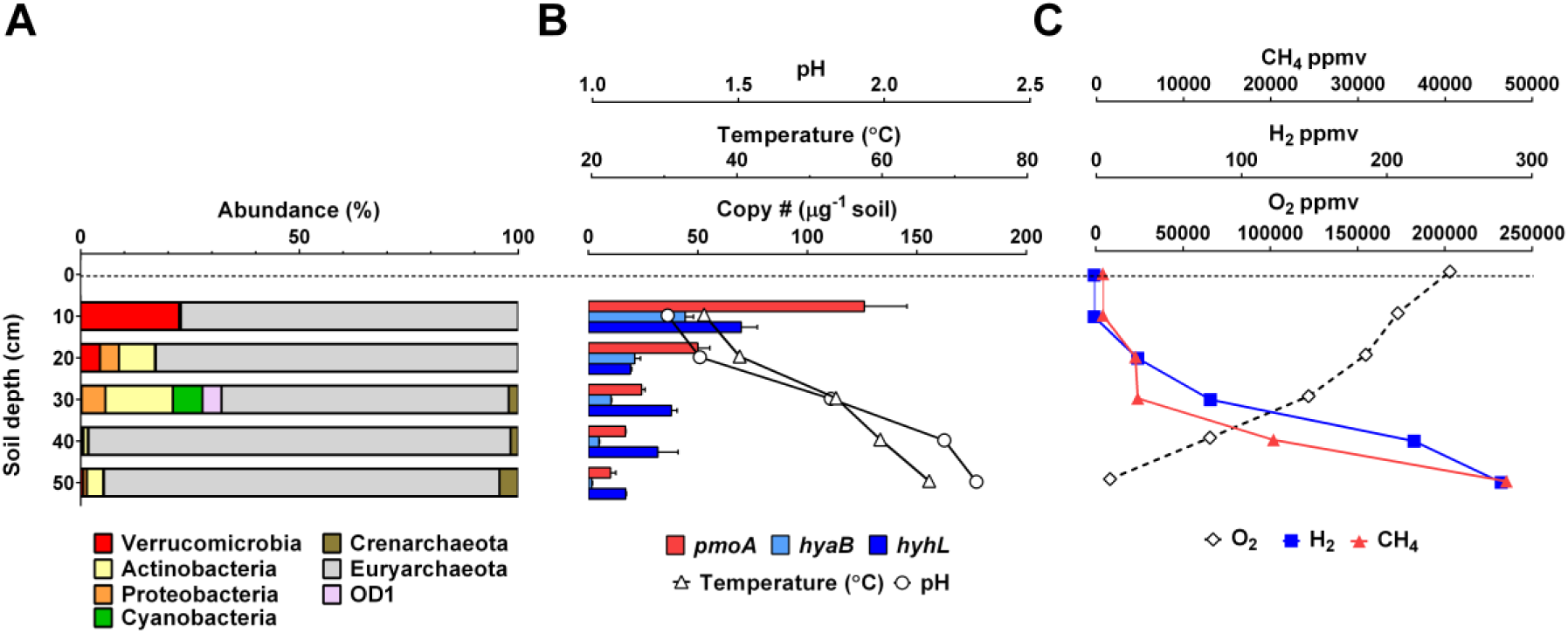
Methanotrophic bacteria are highly abundant in the uppermost soils in Rotokawa, New Zealand. OTUs classified belonging to phylum Verrucomicrobia were the dominant bacterial taxa identified in the top 10 cm. Consistent with a methanotrophic lifestyle, all verrucomicrobial OTUs were further classified into the family Methylacidiphilaceae. 16S rRNA gene sequencing was performed (illumina) on total DNA extracted at 10-50 cm soil depth. Non-rarefied abundance results (%) are shown for all OTUs (> 100 reads) from 130,289 total sequence reads (with an average of 26,058 per sample depth). The abundance (B) of the verrucomicrobial particulate methane monooxygenase (*pmoA*), Group 1d (*hyaB*) and Group 3b (*hyhL*) hydrogenase genes and corresponding (C) soil gas, temperature and pH profiles suggests both high levels of aerobic methane and hydrogen oxidation activity by these acidophilic taxa in the uppermost 20 cm. Error bars shown represent the standard deviation of triplicate measurements.

***Methylacidiphilum* sp. RTK17.1 encodes membrane-bound and cytosolic [NiFe] hydrogenases**. To gain further insight into the hydrogen-oxidising capacity of the Methylacidiphilaceae within the geothermal soil profile, we isolated, characterised, and sequenced a thermotolerant *Methylacidiphilum* sp. (strain RTK17.1; Table S2) from the Rotokawa soils. Characterisation of *Methylacidiphilum* sp. RTK17.1 showed that it optimally grew at pH 2.5 and 50 ºC (T_max_ 58 ºC) and could oxidise methane and fix carbon dioxide in common with the reported observations other *Methylacidiphilum* spp. (12). Glycogen accumulation, as previously described in this genus (33), was also observed (data not shown). Analysis of the *Methylacidiphilum* sp. RTK17.1 genome confirmed the basis for these processes (Table S2) and indicated that the metabolic strategy of this strain is consistent with the other verrucomicrobial methanotrophs. Amperometric assays were performed and demonstrated that the isolate oxidised hydrogen gas, providing the first confirmation that *Methylacidiphilum* spp., and indeed members of the dominant soil phylum Verrucomicrobia can metabolise hydrogen. The microorganism rapidly oxidised hydrogen at rates proportional to cell density (Figure 2a) and consumed hydrogen at concentrations as low as 55 ppmv (Figure S3). As with the closely related *M. infernorum* V4, we noted the presence of two gene clusters encoding [NiFe] hydrogenases in the sequenced genome (Table S2). These enzymes were classified into Groups 1d (*hyaABC*) and 3b (*hyhBGSL*) based on the criteria of our recent survey on environmental hydrogenase distribution (Figure S1) (17). Group 1d enzymes are membrane-bound, hydrogen-uptake [NiFe] hydrogenases that are believed to yield electrons for aerobic respiration via quinone carriers. In comparison, the identified Group 3b [NiFe] hydrogenase belongs to a class of cytosolic enzymes that couples the oxidation of NADPH to the production of hydrogen (34). Reverse transcription (RT) PCR confirmed *Methylacidiphilum* sp. RTK17.1 constitutively expresses both hydrogenases during methanotrophic growth in the presence of hydrogen (Figure S4).

**Figure 2:**
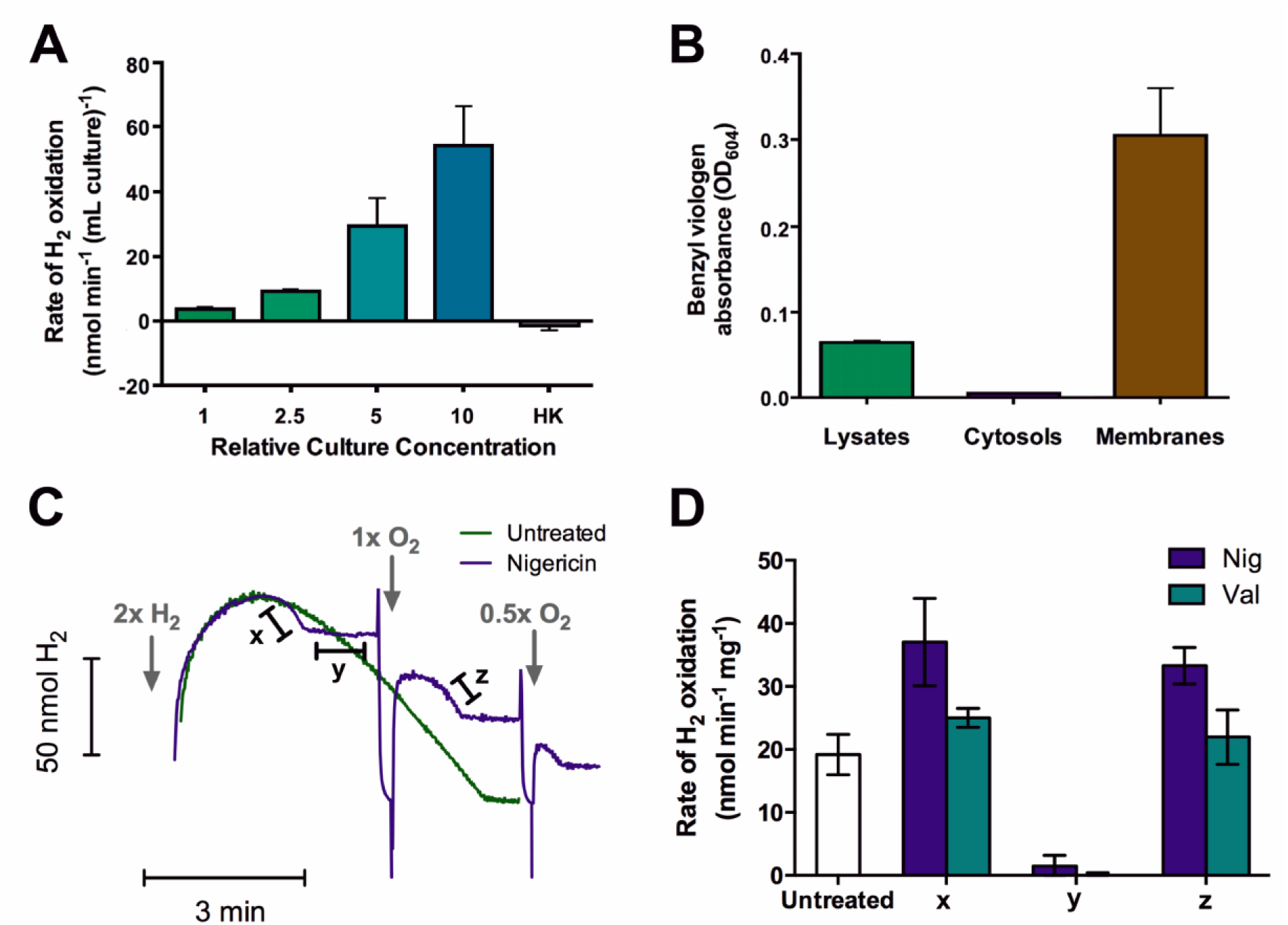
Hydrogenase activity of *Methylacidiphilum* sp. RTK17.1. (A) Whole-cell oxidation of hydrogen measured amperometrically using a hydrogen microsensor. The density-dependence of this process and its sensitivity to heat inactivation (HK) is shown. (B) Colourimetric detection of hydrogenase activity in protein concentration-normalized cell lysates, cytosols, and membranes anaerobically treated with the artificial electron acceptor benzyl viologen. (C) Real-time hydrogen oxidation by untreated cells and nigericin-treated cells. The relative amounts of hydrogen and oxygen added at specific time points is shown. (D) Rates of hydrogen oxidation of untreated, nigericin-treated, and valinomycin-treated cells. For the uncoupler-treated cultures, the initial (*x*), oxygen-limited (*y*), and oxygen restored (*z*) rates of oxidation are shown, which correspond to the rates highlighted in panel (C). Endogenous glycogen catabolism likely contributed to oxygen-limitation (y) observed in nigericin treated cells (Figure S5).

***Methylacidiphilum* sp. RTK17.1 oxidises hydrogen in an aerobic respiratory-linked process**. Cell lysates targeting the Group 1d hydrogenase efficiently coupled the oxidation of hydrogen to the reduction of the artificial electron acceptor benzyl viologen (*E*’_o_ = −374 mV). This activity was localized in the membrane, as shown by the 31-fold increase in benzyl viologen reduction of the membrane fraction relative to the cytosol fraction (Figure 2b). The *hyaC* subunit of this enzyme encodes a predicted membrane-bound *b*-type cytochrome domain known to mediate electron transfer from hydrogen to quinone (35). To probe the physiological interactions of this hydrogenase, we measured real-time hydrogen consumption within cell suspensions. Hydrogen oxidation was stimulated by treatment with the ionophores nigericin and valinomycin respectively (Figures 2c & 2d), showing hydrogenase activity is sensitive to the magnitude of the proton and charge gradients of the proton-motive force. Consistent with being linked to the aerobic respiratory chain, such uncoupled hydrogen oxidation ceased rapidly and could be rescued by supplementation with the terminal electron acceptor, O2; under these conditions the onset of O_2_-limitation was likely exacerbated by the catabolism of endogenous glycogen reserves (Figure S5). By contrast, rates of hydrogen oxidation were reduced following treatment with the protonophore carbonyl cyanide *m*-chlorophenyl hydrazone (CCCP), likely a consequence of secondary intracellular acidification (Figure S6). The Group 1d [NiFe] hydrogenase mediating this process appears oxygen-tolerant, a finding consistent with the sequenced small subunit gene encoding a 6Cys[4Fe3S] cluster, which rapidly reactivates hydrogenases following oxygen inhibition (36, 37). Collectively, these findings demonstrate that this hydrogenase is a strictly membrane-bound, respiratory-linked, oxygen-dependent enzyme that drives ATP synthesis through the Knallgas reaction, i.e. by chemiosmotically coupling hydrogen oxidation to oxygen reduction.

**Oxidation of hydrogen enhances mixotrophic growth**. Pure culture experiments with *Methylacidiphilum* sp. RTK17.1 were performed to determine the physiological roles of the Group 1d hydrogenase. We observed that methane and hydrogen were co-oxidised in both batch bioreactor cultures (Figure S7) and chemostat experiments (Table 1). In these experiments, an exogenous energy source (i.e. hydrogen or methane) was always supplied in excess to suppress the induction of endogenous metabolism and prevent glycogen catabolism (38). Chemostat experiments revealed that hydrogen oxidation is stimulated under hypoxic growth conditions. Observed rates of hydrogen oxidation were 77-fold greater when cells were oxygen-limited compared to oxygen excess conditions (Table 1). Further, under oxygen-limiting conditions with 1.9 % hydrogen addition, the specific consumption rate of hydrogen was 42 % greater than observed rates of methane oxidation. Hydrogen addition into the feedgas significantly (*p*-value < 0.05) increased growth yields (gCDW molCH_4_^-1^) of *Methylacidiphilum* sp. RTK17.1 by 33 % under oxygen excess and 51 % under oxygen-limited conditions (Table 1). Additionally, cellular growth yields (as determined by total protein) of *Methylacidiphilum* sp. RTK17.1, following one week incubation in the presence of 1 % hydrogen and 10 % methane (v/v), were significantly greater than when grown exclusively on methane in batch experiments (Figure S3b). Hydrogen oxidation was also observed both in methane-starved cultures of *Methylacidiphilum* sp. RTK17.1 and following the addition of 4 % (v/v) acetylene (Figure S3a), an inhibitor of methane monooxygenase activity (39). Hence, while hydrogen oxidation alone cannot sustain growth of this methanotroph, it appears sufficient to maintain both the proton-motive force and ATP synthesis required for long-term survival in the absence of the methylotrophic growth substrates.

**Table 1:** Hydrogen oxidation by *Methylacidiphilum sp.* RTK17.1 during chemostat cultivation.

| Condition <sup>a,b</sup> | Growth rate<br>(mg l<sup>-1</sup> h<sup>-1</sup>) | H<sub>2</sub> consumption |  | CH<sub>4</sub> consumption |  | Biomass Yield<sup>d</sup><br>(g<sub>CDW</sub> mol<sub>CH<sub>4</sub></sub><sup>-1</sup>) |
| --- | --- | --- | --- | --- | --- | --- |
|  |  | (mmol l<sup>-1</sup> h<sup>-1</sup>) | (mmol g<sub>CDW</sub><sup>-1</sup> h<sup>-1</sup>) | (mmol l<sup>-1</sup> h<sup>-1</sup>) | (mmol g<sub>CDW</sub><sup>-1</sup> h<sup>-1</sup>) |  |
| O<sub>2</sub> excess, medium H<sub>2</sub> | 9.33 (± 0.52) | 0.01 (± 0.01) | 0.01 (± 0.03) | 1.06 (± 0.12) | 2.40 (± 0.38) | 8.90<sup>‡</sup> (± 1.51) |
| O<sub>2</sub> excess, no H<sub>2</sub> | 8.17 (± 0.82) | 0.00 | 0.00 | 1.22 (± 0.05) | 2.85 (± 0.18) | 6.68 (± 0.41) |
| O<sub>2</sub>-limiting, high H<sub>2</sub> | 5.57 (± 0.16) | 0.78 (± 0.02) | 2.66 (± 0.11) | 0.55 (± 0.03) | 1.89 (± 0.11) | 10.11<sup>¥</sup> (± 0.64) |
| O<sub>2</sub>-limiting, medium H<sub>2</sub> | 4.99 (± 0.17) | 0.33 (± 0.01) | 1.34 (± 0.06) | 0.70 (± 0.02) | 2.80 (± 0.13) | 7.16 (± 0.34) |
| O<sub>2</sub>-limiting, low H<sub>2</sub> | 5.60 (± 0.31) | 0.19 (± 0.00) | 0.73 (± 0.04) | 0.71 (± 0.01) | 2.68 (± 0.16) | 7.87<sup>¥</sup> (± 0.45) |
| O<sub>2</sub>-limiting, no H<sub>2</sub> | 5.40 (± 0.26) | 0.00 | 0.00 | 0.80 (± 0.05) | 3.11 (± 0.27) | 6.79 (± 0.55) |

<sup>a</sup> Feedgas was continuously supplied at a rate of 10 ml min<sup>-1</sup> with the following composition % (v/v): O<sub>2</sub> Excess, 14.1 %; O<sub>2</sub> limiting, 3.5 %. High, medium and low H<sub>2</sub> experiments consisted of 1.9 %, 0.7 % and 0.4 % H<sub>2</sub> supply. For all experiments excess CH<sub>4</sub> was supplied at 3 %, CO<sub>2</sub> at 26 % with the balance made up with N<sub>2</sub>.
<sup>b</sup> a dilution rate of 0.02 h<sup>-1</sup> was maintained for all experiments
<sup>c</sup> gram cell dry weight (g<sub>CDW</sub>) measurements were determined by correlating OD<sub>600</sub> measurements to an existing standard curve (OD<sub>600</sub> = 1 corresponds to 0.43 g<sub>CDW</sub>/l).
<sup>d</sup> Standard deviation shown in brackets are calculated from a minimum of triplicate measurements
<sup>¥</sup> denotes significant increase (*p*-value < 0.05) in yield from O<sub>2</sub>-limiting, no H<sub>2</sub> result; unpaired *t*-test, one tailed.
<sup>‡</sup> denotes significant increase (*p*-value < 0.05) in yield from O<sub>2</sub> excess, no H<sub>2</sub> result; unpaired *t*-test, one tailed.

## Discussion

This study shows a methanotrophic bacterium is able to constitutively consume methane and hydrogen, either together or separately depending on gas availability, to meet energy needs. We demonstrated that the genetic determinants of aerobic hydrogen respiration, specifically genes encoding a member of the Group 1d [NiFe] hydrogenases, are encoded and expressed in the verrucomicrobial methanotroph *Methylacidiphilum* sp. RTK17.1. The strain rapidly oxidised exogenous hydrogen with this enzyme in a membrane-bound aerobic respiratory process and likely provides hydrogen-derived reducing equivalents to the quinone pool for methane monooxygenase activity and/or cytochrome reduction, *via* activity of a recently described alternative (ACIII) cytochrome complex (40). Our model of how methane, methanol and hydrogen oxidation are integrated into the respiratory chain of verrucomicrobial methanotrophs is shown in Figure 3. Through this metabolic flexibility, we propose that *Methylacidiphilum* sp. RTK17.1 is able to balance the catabolic and anabolic requirements for reducing energy imposed by its challenging methanotrophic lifestyle and the methane monooxygenase reaction. This flexibility likely contributes to the dominance of the Methylacidiphilaceae in aerated acidophilic geothermal soils at Rotokawa, New Zealand. As a dominant genus in the uppermost 20 cm of these soils, *Methylacidiphilum* sp. RTK17.1 can simultaneously oxidise geothermally-produced methane and hydrogen gases while consuming atmospheric oxygen and fixing carbon dioxide and nitrogen gases. In addition to conferring a survival advantage during periods of methane limitation, our data shows hydrogen oxidation may be favoured during hypoxia. When grown under oxygen-limiting conditions, rates of hydrogen consumption in cultures of *Methylacidiphilum* sp. RTK17.1 increased by 77-fold and exceeded observed rates of methane oxidation (Table 1). Similarly, a transcriptome study investigating nitrogen fixation reported that the Group 1d hydrogenase in *Methylacidiphilum fumarolicum* SolV was upregulated 20-fold in response to oxygen limitation (41). This preference for hydrogen suggests methanotrophic bacteria occupying the soil oxic/anoxic interface display a predominately mixotrophic lifestyle. In fact, under oxygen limited experimental conditions, up to 32 % of the total reducing energy supplied to the respiratory chain may be provided from hydrogen-derived electrons. Recalling that pMMO requires oxygen as a substrate to catalyse the conversion of methane to methanol, mixotrophy through hydrogen respiration both sustains the energy requirements for cell growth while reducing the overall demand for oxygen. Given the energetic requirement for cell maintenance is three orders of magnitude less than for growth (42, 43), it is probable that hydrogen oxidation serves as a reliable mechanism for the persistence of methanotrophic bacteria in these habitats.

**Figure 3:**
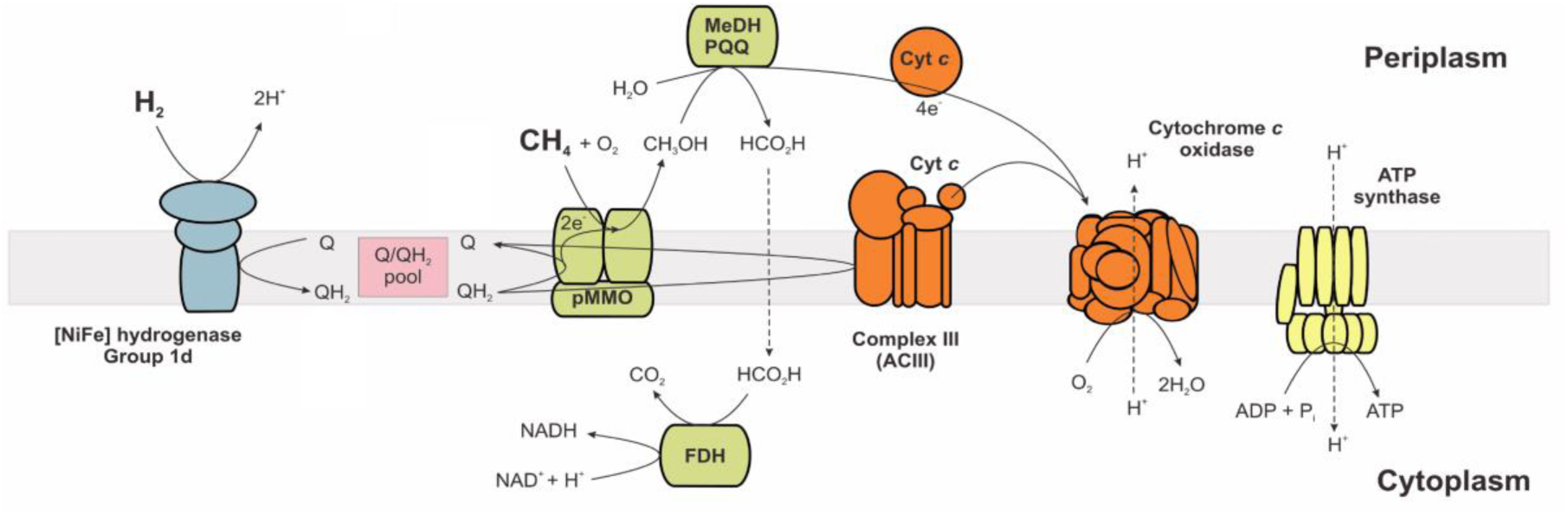
Proposed model of respiratory-linked methane and hydrogen oxidation in *Methylacidiphilum* sp. RTK17.1. During mixotrophic growth, the oxidation of both hydrogen (H_2_) and methane (CH_4_) yields reducing equivalents (QH_2_) for the respiratory chain, a large proton-motive force is generated, and sufficient ATP is produced for growth via an H^+^-translocating F_1_F_o_-ATP synthase. Some of the quinol generated through H_2_ oxidation provides the electrons necessary for pMMO catalysis. Following CH_4_ oxidation by pMMO, ensuing reactions catalysed by an XoxF type methanol dehydrogenase (MeDH) and formate dehydrogenase (FDH) contribute additional reductant (Cyt *c* and NADH) into the respiratory chain for ATP production and growth (11). In methane-limiting conditions, the identified Group 1d [NiFe] hydrogenase generates a small proton-motive force, oxidised reductant (Q) is maintained via type ACIII cytochrome complex (40) and sufficient ATP is generated for maintenance. Respiratory complexes I and II are not shown but are encoded in the genome of *Methylacidiphilum* sp. RTK17.1 (Table S2).

Genomic evidence supports that hydrogen oxidation is a commonly utilised strategy for the growth and survival of methanotrophic bacteria. Hydrogenases were identified in all 31 genomes of methanotrophic bacteria surveyed (Figure S1). Previous studies have shown two model methanotrophs, *Methylosinus trichsporium* OB3b (21) and *Methylococcus capsulatus* (Bath) (24), oxidise hydrogen but the physiological role of their encoded hydrogenases has never been fully investigated. Consistent with hydrogen contributing to the persistence of methanotrophic bacteria, genes encoding the Group 1h [NiFe] hydrogenases, while absent from *Methylacidiphilum* sp. RTK17.1, were identified in the genomes of selected alphaproteobacterial and verrucomicrobial methanotrophs. These high-affinity enzymes are capable of oxidising tropospheric concentrations of hydrogen and have been linked to persistence within the dominant soil phyla Actinobacteria (20) and Acidobacteria (18). Our genomic analysis indicates methanotrophic bacteria may additionally evolve hydrogen fermentatively (*via* Group 3b and 3d [NiFe] hydrogenases), which by analogy with *Mycobacterium smegmatis*, may be important for surviving anoxia (34). They also encode the capacity to sense hydrogen (*via* Group 2b and 2c [NiFe] hydrogenases) which may serve to regulate expression of uptake hydrogenases and also potentially global cellular processes. Hydrogen is likely an attractive energy source for these microorganisms because of its high-diffusibility, relative dependability, and high energy content. Whereas biogenic methane is exclusively produced by methanogenic archaea, hydrogen is biologically produced by diverse organisms across the three domains of life as a result of fermentation, photobiological processes, and nitrogen fixation (17). Hence, the ability of methanotrophic bacteria to utilise this widespread energy source may enhance their capacity to colonise environments and adapt to environmental change. This hidden metabolic flexibility may explain why the range of low-affinity methanotrophic bacteria extends beyond ecosystems with high methane fluxes (44).

By demonstrating that respiratory-linked hydrogen oxidation enhances growth, we reveal that verrucomicrobial methanotrophs are mixotrophs capable of growth and survival on hydrocarbon and non-hydrocarbon substrates alike. The narrative that methanotrophic bacteria are niche specialists, with obligately methylotrophic lifestyles, is based on studies under optimal growth conditions and ignores the requirement of these organisms to adapt to methane and/or oxygen limitation. Particularly within dynamic oxic/anoxic boundaries, where methanotrophic bacteria occur in high abundance (45), a specialist metabolism is likely to reduce the overall fitness of these microorganisms. We therefore propose that mixotrophic growth and survival through hydrogen metabolism provides a general mechanism by which this contradiction is resolved. Expanding the metabolic flexibility of methanotrophs will have broad ecological implications and result in an improved understanding of global methane and hydrogen cycles.

## Methods

**Environmental sampling.** Soil samples (~ 50 g) were collected every 10 cm from the surface of the sampling site (38º37’30.8”S, 176º11’55.3”E) to a maximum depth of 50 cm. This site, located within Rotokawa geothermal field was selected on the basis of previous research showing elevated soil gas concentrations (25). Soil temperatures were measured in the field using a 51II single input digital thermometer (Fluke). The pH was measured upon returning to the laboratory using a model HI11310 pH probe (Hanna Instruments). Soil gas samples were collected every 10 cm using a custom built gas-sampling probe equipped with a 1 | gas tight syringe (SGE Analytical Science). Gas samples were collected and stored at 25 ºC in 50 ml Air & Gas Sampling Bags (Calibrated Instruments) and were processed within 48 h on a 490 MicroGC (Agilent Technologies, Santa Clara, CA) equipped with Molecular Sieve 5A with heated injector (50 °C, back-flush at 5.10 s, column at 90 °C, 150 kPa), a PoraPak Q column with heated injector (50 °C, no back-flush, column at 70 °C,150 kPa) and a 5CB column with heated injector (50 °C, no back-flush, column at 80 °C, 150 kPa). Methane and hydrogen gas consumption by soil microbial communities was determined by incubating 1 g of soils collected from the Rotokawa sampling site (depth < 10 cm) in 112 ml gas-tight serum bottles at 37 and 50 ºC. The serum bottle headspace were air emended with 300 ppmv and 400 ppmv of hydrogen and methane gas respectively. Headspace methane and hydrogen gas concentrations were measured with a PeakPerformer gas chromatograph equipped with a flame ionising detector (FID: methane) and a PP1 Gas Analyzer equipped with a reducing compound photometer (RCP: hydrogen).

**Isolation and cultivation of *Methylacidiphilum* sp. RTK17.1.** Soil samples (1 g) were inoculated into serum bottles containing 50 ml media (pH 2.5). All cultivations were performed in a V4 mineral medium as described previously (29) but with the addition of rare earth elements lanthanum and cerium (27). Methane (10 % v/v) and carbon dioxide (1 %) were added to an air headspace and samples were incubated at 60 ºC with shaking 150 rpm. Methane in the headspace was monitored with a PeakPerformer gas chromatograph equipped with a flame ionising detector (FID). Following several passages (10 % v/v) into liquid media, enrichments were transferred onto solid media. Following several weeks incubation (60 ºC) single colonies were re-streaked before being transferred back into liquid media. Isolate identity was confirmed via sequencing of the 16S rRNA gene (Macrogen) using bacterial 9f/1492r primers (46).

### Chemostat cultivation and batch experiments

Chemostat cultivation of *Methylacidiphilum* sp. RTK17.1 was performed to investigate the influence of hydrogen on growth under oxygen excess and oxygen-limiting conditions. A 1 l bioreactor (BioFlo 110, New Brunswick Scientific. Edison, NJ, USA) was used for these studies. Cultures were incubated at 50 ºC and pH 3 with continuous stirring (800 rpm). Bioreactor volume was kept constant at 0.5 l by automatic regulation of the culture level. V4 mineral media was supplied at a constant flow rate of 10 ml h^-1^ (D=0.02 h^-1^). Dissolved oxygen was monitored using a InPro 6810 Polarographic Oxygen Sensor (Mettler-Toledo). Custom gas mixtures were prepared in a compressed gas cylinder and supplied to the chemostat at a rate of 10 ml min^-1^ using a mass flow controller (EL-FLOW, Bronkhorst, Netherlands). Gas mixtures contained approximately (v/v) 3 % methane and 26 % carbon dioxide for all experiments. For oxygen excess and oxygen-limiting conditions, (v/v) 14.1 % and 3.5 % oxygen was supplied. High, medium and low hydrogen experiments consisted of (v/v) 1.9 %, 0.7 % and 0.4 % hydrogen additions. The balance of all gas mixtures was made up with nitrogen.

Cell density in liquid samples was monitored by measuring turbidity at 600 nm using a Ultrospec 10 cell density meter (Amersham Bioscience). One unit of OD_600_ was found to be equivalent to 0.43 g l^−1^ cell dry weight (CDW) for *Methylacidiphilum* sp. RTK17.1. After achieving a steady state condition as determined by OD600, influent and effluent gas concentrations were monitored over several days using a 490 MicroGC (Agilent Technologies, Santa Clara, CA). Biomass cell dry weight was used to calculate growth rate and specific gas consumption rate.

To investigate whether *Methylacidiphilum* sp. RTK17.1 oxidises ecologically significant concentrations of hydrogen, 350 ml cultures (in triplicate) were incubated in 1 l rubber-stoppered Schott bottles in a headspace of air supplemented with 100-250 ppmv hydrogen, 1 % (v/v) methane and 1 % carbon dioxide. Acetylene gas was added in some experiments (4 % v/v) to inhibit MMO activity as previously described (39). Finally, to determine if growth was enhanced in the presence of hydrogen, 20 paired cultures (*n* = 40) of *Methylacidiphilum* sp. RTK17.1 were incubated with or without 1 % (v/v) hydrogen in an air headspace supplemented with (v/v) 10 % methane and 1 % carbon dioxide. Cultures were incubated in a custom Test Tube Oscillator (TTO; agitation 1.2 Hz; Terratec). Following seven days incubation, total protein was determined as described previously (47). Statistical significance of observed differences of growth yields was determined using a *student’s t-test* (α = 0.05). Headspace mixing ratios of hydrogen and methane were monitored throughout batch experiments as described above.

**Hydrogenase activity measurements.** Hydrogenase activity of *Methylacidiphilum* sp. RTK17.1 was measured in stationary-phase cultures harvested from the bioreactor. For amperometric measurements, whole-cells were concentrated 5-, 10-, 20-, and 30-fold by centrifugation followed by resuspension in phosphate-buffered saline (pH 7.4). Rate of hydrogen oxidation was measured at 50 °C using a hydrogen-MR hydrogen microsensor (Unisense) as previously described (18, 48). For colourimetric assays, 500 ml culture was harvested by centrifugation (15 min, 5,000 × g, 4 °C) and treated as previously described (18, 20) to prepare crude, cytosolic and membrane fractions for analysis. To test for hydrogenase activity, samples (20 µg protein) from each cell fraction were incubated with 1 ml 50 mM potassium phosphate buffer (pH 7.0) and 50 µM benzyl viologen for eight hours in an anaerobic chamber (5 % hydrogen, 10 % carbon dioxide, 85 % nitrogen (v/v)). Debris was removed by centrifugation (15 min, 10,000 × g, 4 °C) and the absorbance of the supernatants was read at 604 nm in a Jenway 6300 spectrophotometer.

Additional methodologies for nucleic acid extraction, PCR, genome sequencing, oxygen consumption experiments and soil microbial community determination can be found in the supporting information.

## Acknowledgments

This work was supported by Geothermal Resources of New Zealand (CRC, MBS, KMH, JFP) and Environmental Technologies (CRC, DG, BM, CC) research programmes at GNS Science and Scion respectively. CRC was further supported by Genomes Canada through the Applied Genomics Research in BioProducts or Crops (ABC) program for the Grant titled “Microbial Genomics for Biofuels and Co-Products from Biorefining Processes” and by the government of the Province of Manitoba through the Manitoba Research Innovation Fund (MIRF). CG was supported by a CSIRO Office of the Chief Executive Postdoctoral Fellowship and KH was supported by a University of Otago Postgraduate Scholarship. The authors wish to thank the Department of Conservation for their assistance in sampling the Rotokawa geothermal site. Relevant gene sequences (accession numbers KU509367-KU509352) have been deposited into GenBank for archival storage.

## Author contributions

CRC, CG, MBS, GC, KMH, KH, CC, DJG, RS and BM contributed to experimental design. Bioreactor and wet lab experiments were conducted by CRC, BM, CC, DJG and KMH. CRC, CG and KH undertook cell respiratory analysis. CRC and MBS performed fieldwork and genomic analyses. JFP performed the microbial community analysis. CRC, MBS, CG, KH and GC wrote the manuscript.

## Supporting Information

### Supplementary Methods

#### Nucleic acid extraction

*Methylacidiphilum* sp. RTK17.1 genomic DNA was extracted using the NucleoSpin Tissue kit (Macherey-Nagel) as per the manufacturer’s instructions for difficult to lyse Gram-positive bacteria. DNA was extracted from Rotokawa soil samples using the NucleoSpin Soil DNA extraction kit (Macherey-Nagel) as per the manufacturers recommended protocol.

For RNA isolation from *Methylacidiphilum* sp. RTK17.1, stationary phase (Abs_600nm_ = 4.0, 10 ml cultures) cells were harvested by centrifugation (27,000 × g, 20 min, 4 °C), resuspended in 1 ml RNA Later (ThermoFisher Scientific) and stored at −20 ºC as recommended. Cell lysis, total RNA extractions and cDNA synthesis were performed as previously described (18).

#### PCR, RT-PCR, Quantitative PCR and genome sequencing

*Methylacidiphilum* sp. RTK17.1 genomic DNA and cDNA (10 ng) was (RT)PCR amplified using i-Taq DNA polymerase (2.5 U; Intron Biotechnology) in 50 µl reactions containing dNTPs (1 mM), 1X PCR Buffer, MgCl_2_ (1.5 mM), forward primer (0.5 µM) and reverse primer (0.5 µM). Amplifications conditions using an Mx3000p thermocycler (Strategene) were as follows; an initial 94 ºC for 5 min; followed by 30 cycles of 94 ºC for 45 s; 55 ºC for 45 s; 72 ºC for 90 s; and a final extension at 72 ºC for 5 min. PCRs were run through electrophoresis on a 1 % (w/v) TAE agarose gel with 1X RedSafe nucleic acid staining solution and visualised on a GeneGenius Bio Imaging System (Syngene).

Quantitative-PCR (qPCR) was performed to investigate the abundance of Verrucomicrobial methanotrophs type by probing for MMO (*pmoA*) and hydrogenase genes (Group 1d and 3b) within the Rotokawa soil profile. Total DNA extracted from 1 g soil was amplified using a Probe Fast Universal qPCR kit (KAPA Biosystems) according to the manufacturer’s instructions. Primer sequences for all qPCR amplifications are provided in Table S2. qPCR reactions were optimised and conducted on an Mx3000p thermocycler (Strategene). Abundance (expressed per µg Soil) of MMO and hydrogenase genes in DNA samples was calculated from cycle threshold values (Ct) and standard curves.

To prepare qPCR standard curves, amplicons were cloned following the recommended protocol for the TOPO-TA Cloning kit (ThermoFisher Scientific). Transformation of competent *Eschericia coli* cells was achieved using a MicroPulse electroporator (BioRad) at setting EC1. Standard curves for qPCR were prepared from the serial dilution of plasmid DNA recovered using the Zippy Plasmid Miniprep kit (Zymo Research) following Qubit fluorometirc quantification (ThermoFisher Scientific).

The genome of *Methylacidiphilum* sp. RTK17.1 was sequenced using the PacBio RS platform (Macrogen). Genome assembly was performed, *de novo*, via the SMRT Portal using hierarchical genome-assembly process (HGAP) pipeline (49). A high quality *de novo* assembly was generated from three SMRT sequencing runs the HGAP.3 protocol. A pre-assembly genome construct was assembled using Celera Assembler and then polished using Quiver (49). Quality assessment of the *de novo* assembly was conducted using BridgeMapper. Gene prediction was performed using Quiver and genome annotation was completed using the integrated microbial genomes database pipeline (50).

#### Soil microbial community determination

The V4 region of the 16S rRNA gene was sequenced using the Illumina MiSeq^®^ System. Sequences were analysed using UPARSE pipeline for quality filtering and clustering (51). Briefly, paired end reads were merged and those less than 200 or 350 bp were discarded, for 515f/806r and 341/785r primer amplicons respectively (Table S1). Quality filtering was applied with a maximum expected error value of 1. Retained sequences were dereplicated and unique sequences removed. Next, reads were clustered to 97 % similarity, which includes a chimera check, and a *de novo* database was created of representative operational taxonomic units (OTUs). Original pre-filtered sequences were mapped to these OTUs, and taxonomy was assigned using the Ribosomal Database Project classifier (with a minimum confidence score of 0.5) (52) against the GreenGenes 16S rRNA database (13_8 release) (53). Final count number was 130,289 reads across all depths for the 515f/806r primer set, with a total of 48 differential OTUs (492,588 for 341/785r, with 107 OTUs).

#### Static liquid fed-batch bioreactor cultivations

*Methylacidiphilum* sp. RTK17.1 was cultivated to stationary phase for subsequent hydrogenase activity and oxygen respiration measurements in a semi-continuous fed-batch bioreactor (New Brunswick; volume 1l, pH control 2.5, temp 50 ºC, agitation 100 rpm) in an artificial headspace composed of 10 % methane, 10 % hydrogen, 2 0 % O_2_, 40 % carbon dioxide (v/v, balance N_2_; flow rate 60 ml min^-1^) equipped with headspace recirculation and automated sampling via a 490 MicroGC (Agilent Technologies, Santa Clara, CA). Gas mixtures were supplied for 2 min every hour at a rate of 60 ml min^-1^.

#### Oxygen consumption experiments

Oxygen consumption experiments were performed on cell suspensions of *Methylacidiphilum* sp. RTK17.1 to determine the influence of endogenous glycogen catabolism on O_2_ dependent hydrogenase measurements. For these experiments, 2 mL cells (OD_600_ 1.0) were added to a Clarke-type oxygen electrode and incubated at 50 ºC for up to 12 min without the addition of exogenous energy sources. Cell suspensions were treated with the protonophore (1 µM) CCCP, 1 mM Iodoacetamide (an inhibitor of glycolysis) and 1 mM potassium cyanide (KCN) to determine whether observed rates oxygen consumption were a consequence of glycogen catabolism.

**Figure S1:**
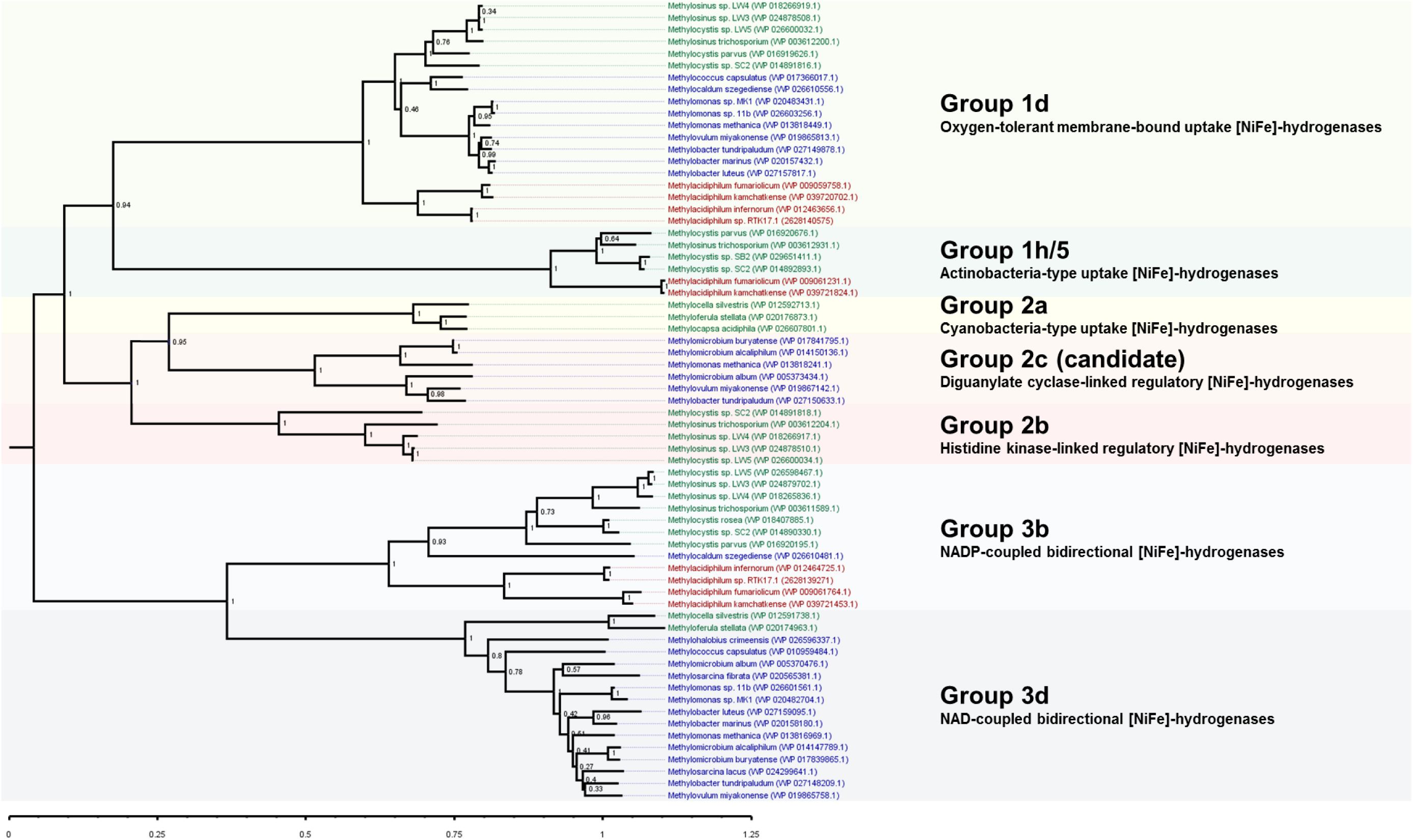
The classification and phylogeny of the [NiFe] hydrogenase large subunit identified in the genomes of methanotrophic bacteria.

**Figure S2:**
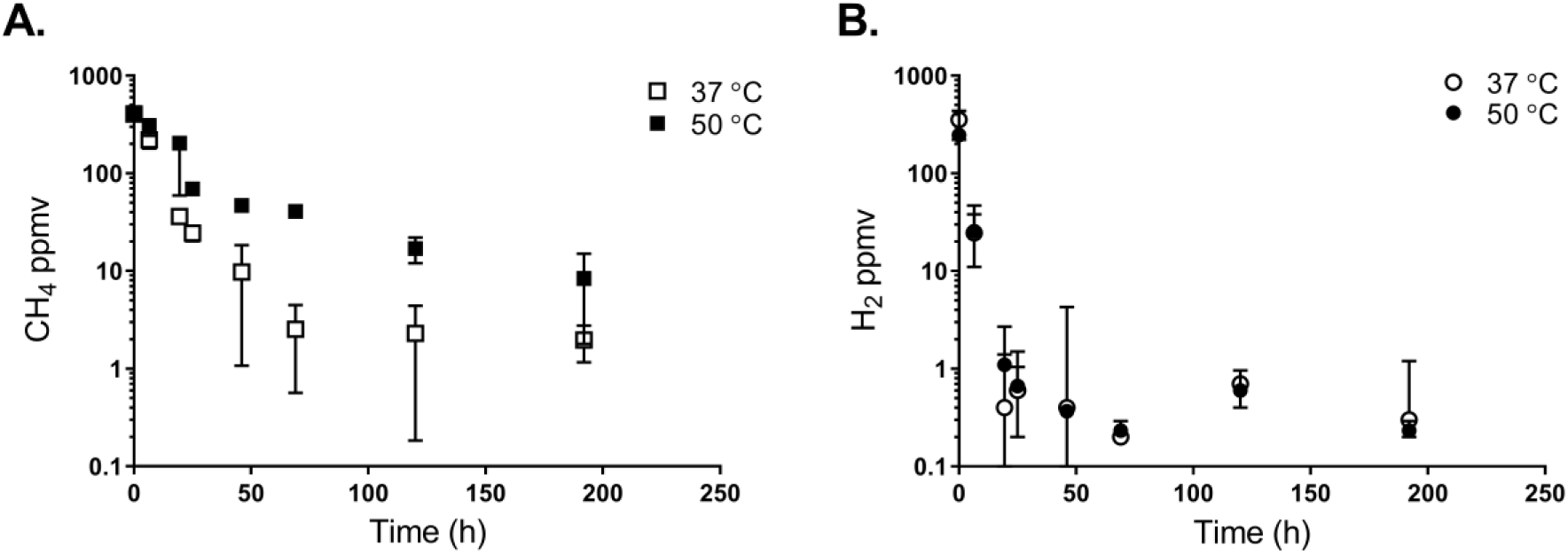
Oxidation of CH_4_ and H_2_ by surface Rotokawa soils (< 10 cm depth) incubated at mesophilic (37 ºC) and thermophilic (50 ºC) temperatures. Soils samples (1 g) were collected and incubated following addition of A) ~400 ppmv CH_4_ and B) ~300 ppmv H_2_ into the headspace. The average and standard deviation of triplicate samples is shown.

**Figure S3:**
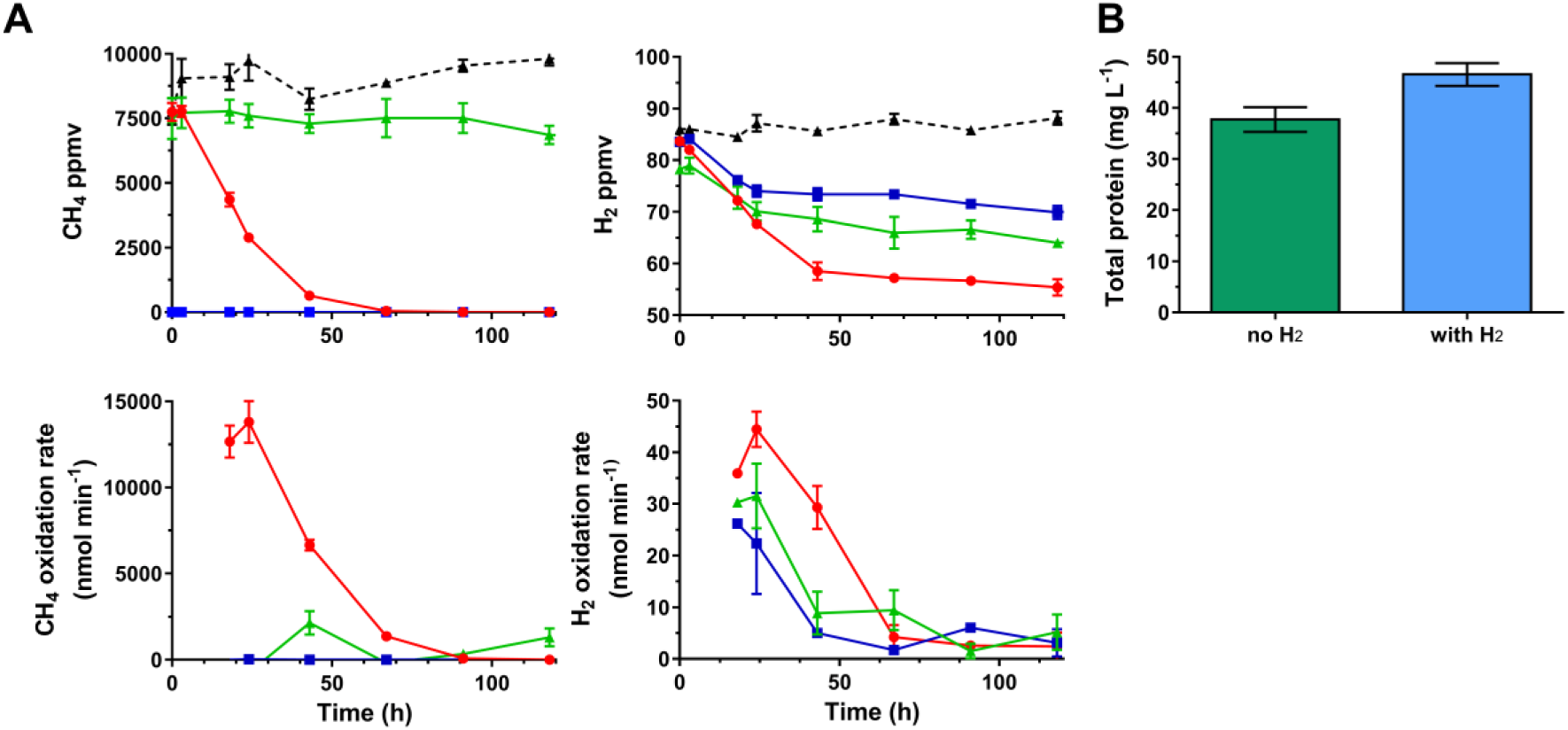
Batch culture growth experiments and observations of methane and hydrogen oxidation for *Methylacidiphilum* sp. RTK17.1 following treatment with acetylene. (A) Rates of hydrogen oxidation are greatest during methanotrophic growth (red circle) but are also measurable in the presence of 4 % (v/v) acetylene gas (an inhibitor of pMMO; green triangle) and in the absence of methane (blue square). Black triangles with dashed lines denote negative controls. Observed rates of methane and hydrogen oxidation decline as substrates are exhausted. Error bars shown represent the standard deviation of biological triplicate samples. (B) Methanotrophic growth of *Methylacidiphilum* sp. RTK17.1 is enhanced by the addition of 1 % H_2_ (v/v). Differences in growth observed (as determined by total protein) with and without hydrogen addition were analysed following 1 week incubation (50 ºC) in 20 paired samples by one-tailed t-test assuming Gaussian distribution and determined to be significant (*p-value* 0.007, *df=* 19, SEM error bars are shown).

**Figure S4:**
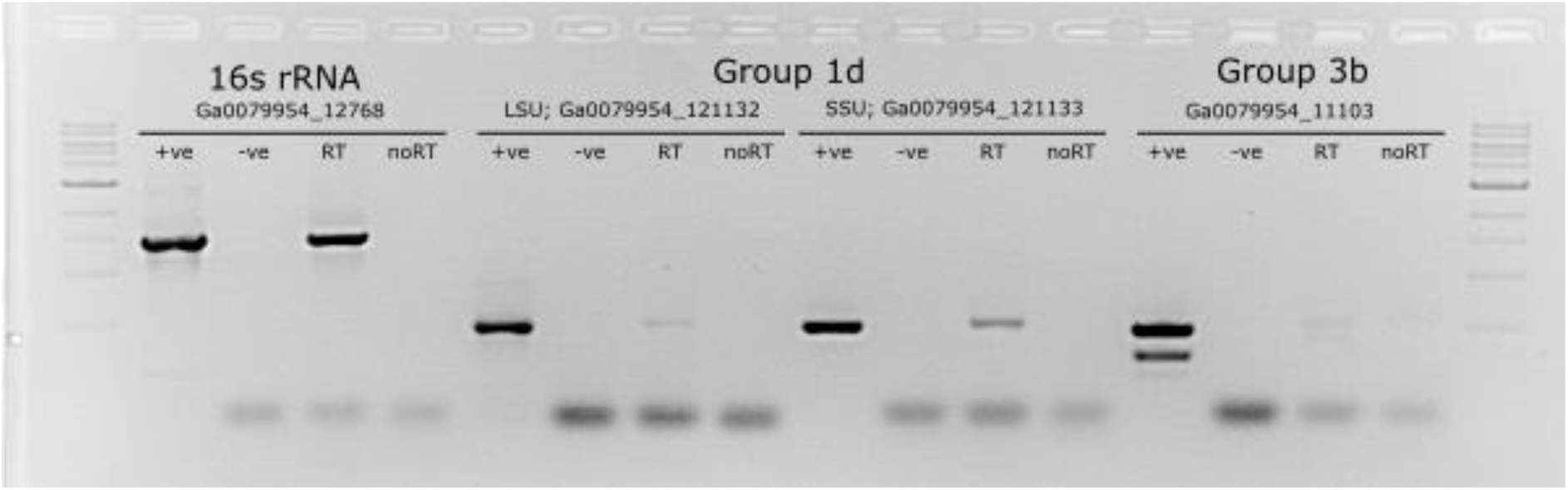
Reverse Transcriptase PCR of total mRNA isolated from stationary phase cultures of *Methylacidiphilum* RTK17.1 confirms the expression of the membrane bound Group 1d [NiFe] hydrogenase. Only faint amplification of cDNA encoding the cytosolic Group 3b enzyme was observed.

**Figure S1:**
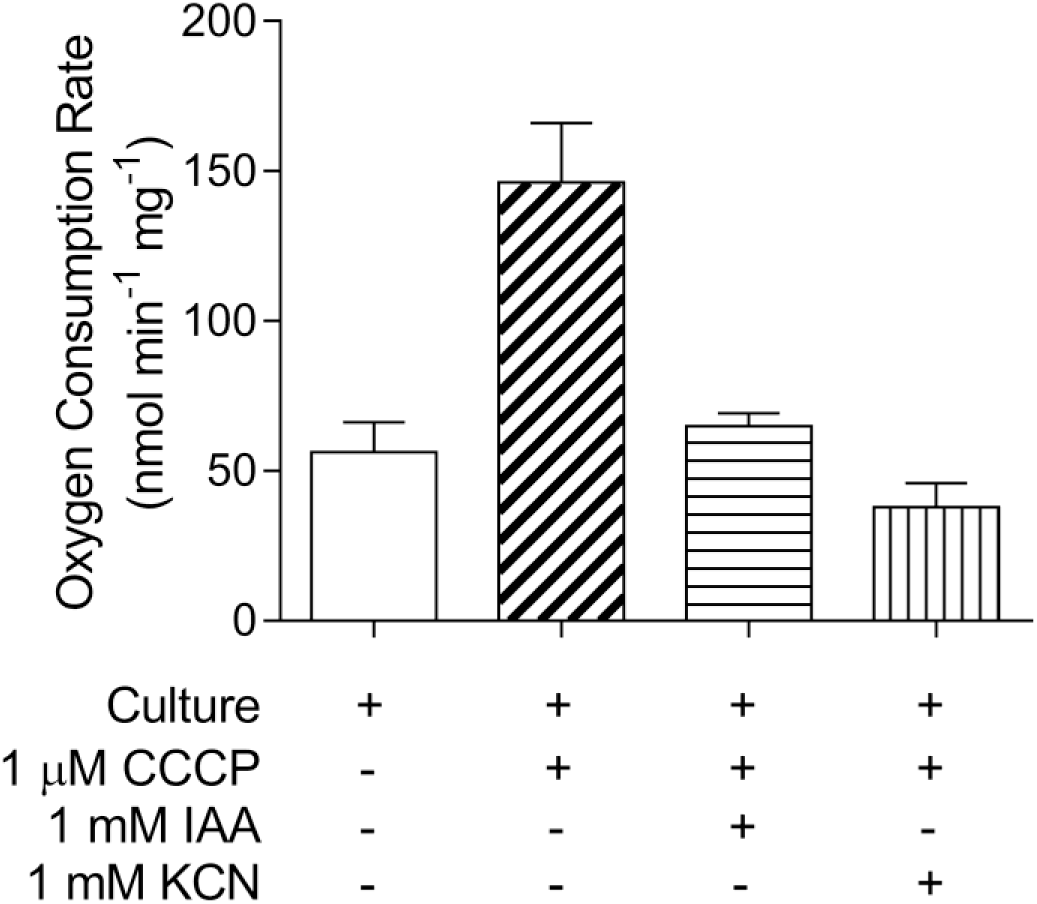
Treatment of cell suspensions of *Methylacidiphilum sp.* RTK17.1 with the uncoupler carbonyl cyanide *m*-chlorophenyl hydrazone (CCCP), the glycolytic inhibitor iodoacetamide (IAA) and the respiratory chain inhibitor potassium cyanide (KCN). This indicates that observed oxygen consumption, in the absence of exogenous energy sources (H_2_ or CH_4_), is likely a consequence of endogenous glycogen catabolism.

**Figure S6:**
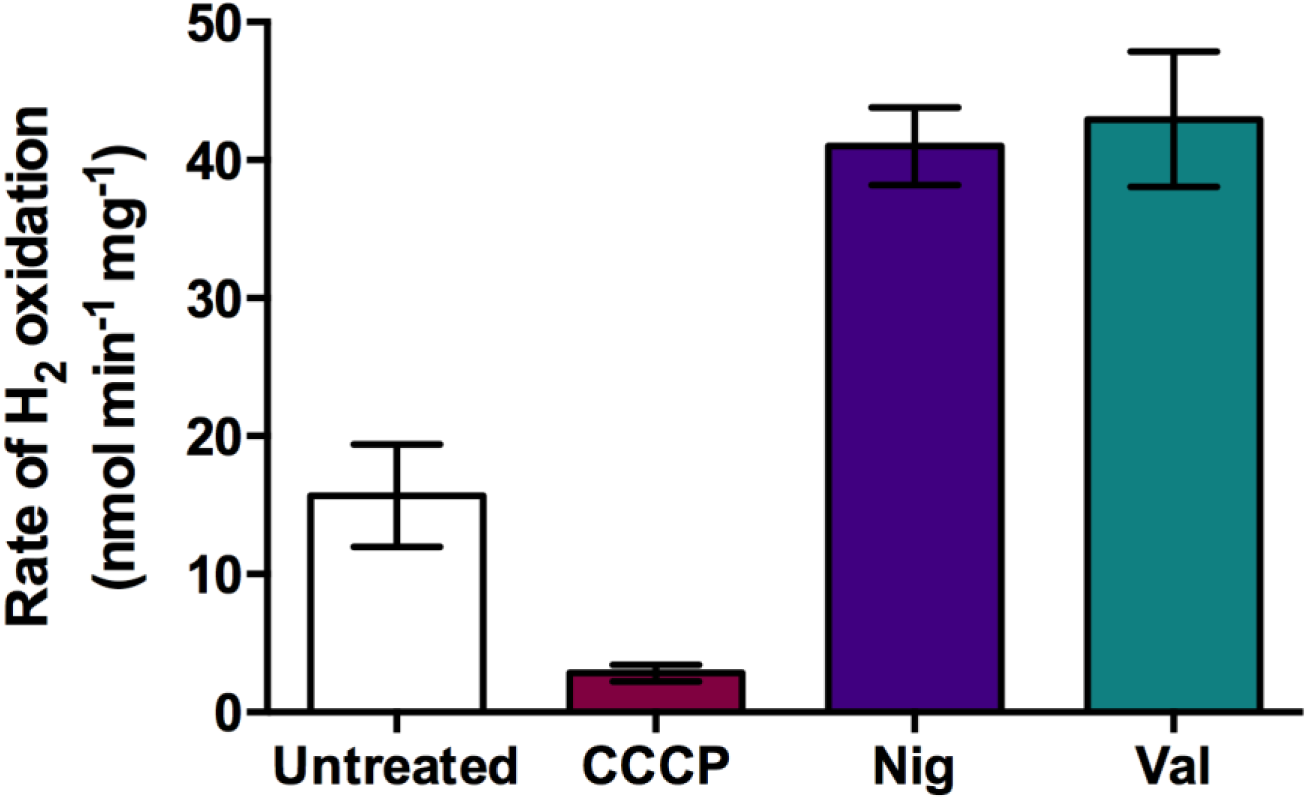
Treatment of concentrated cell suspensions of *Methylacidiphilum sp.* RTK17.1 with the ionophores nigericin (Nig) and valinomycin (Val) indicates hydrogenase activity is sensitive to the magnitude of the proton- and charge-gradients of the proton-motive force. The reduction in rate of hydrogen oxidation observed when treated with carbonyl cyanide *m*-chlorophenyl hydrazone (CCCP) is likely a consequence of secondary intracellular acidification.

**Figure S7:**
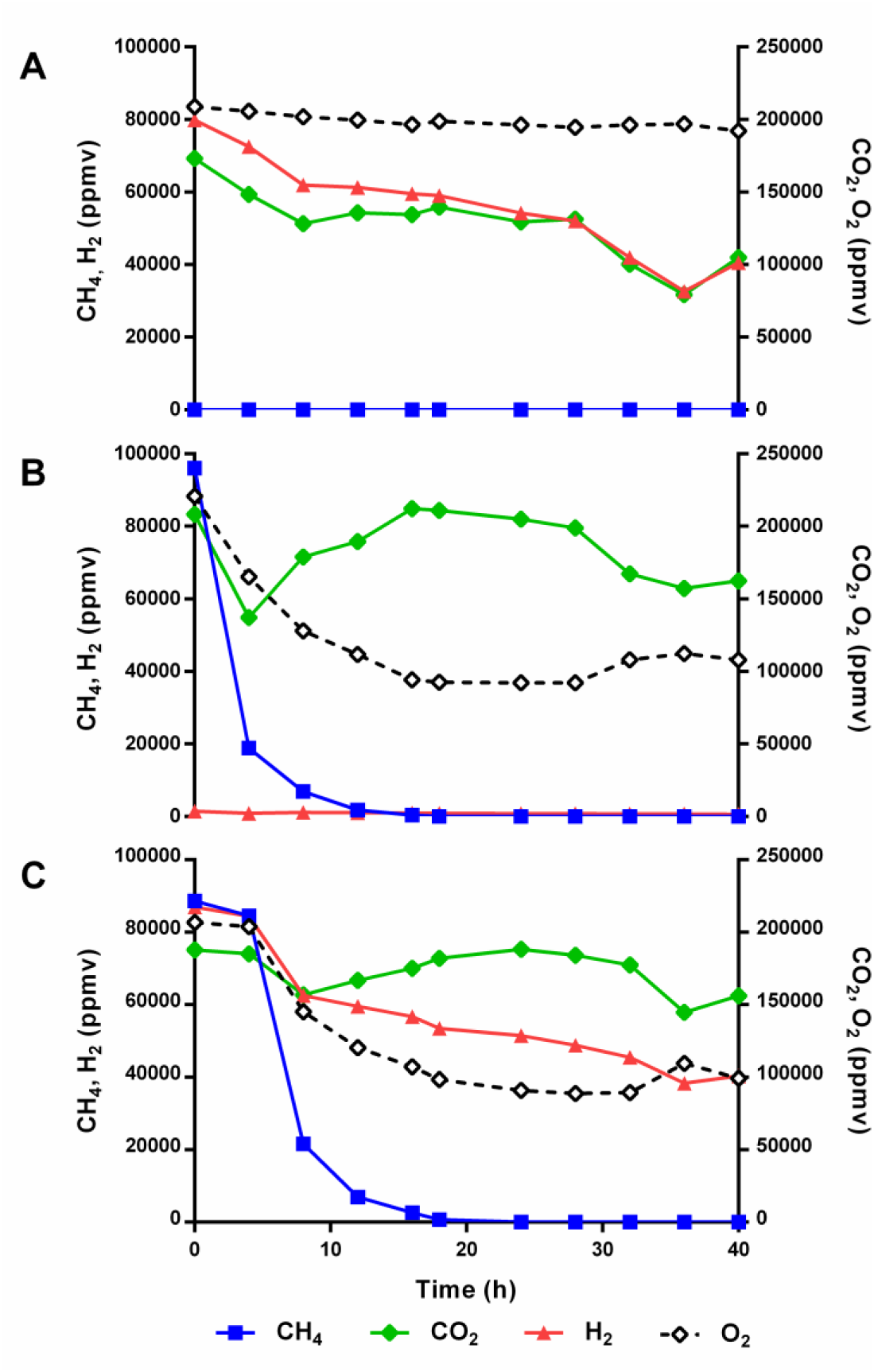
*Methylacidiphilum* sp. RTK17.1 oxidises hydrogen (H_2_) and fixes carbon dioxide (CO_2_) in the absence of methane (CH_4_). In static liquid fed-batch bioreactor experiments, stationary-phase cultures were incubated (50 ºC) in a nitrogen headspace with mixing ratios (v/v) as follows: (A) 0 % CH_4_, 9 % H_2_, 19 % CO_2_, 20 % O_2_; (B): 9 % CH_4_, 0 % H_2_, 19 % CO_2_, 20 % O_2_; (C): 9 % CH_4_, 9 % H_2_, 19 % CO_2_, 20 % O_2_. In the absence of CH_4_ or H_2_ addition, Argon (10 %) supplementation was used to maintain equivalent headspace mixing ratios of the other gases between experiments. We did not determine whether CO_2_ fixation in the absence of CH_4_ was driven by H_2_ oxidation or instead metabolism of endogenous glycogen reserves.

**Table S1:**
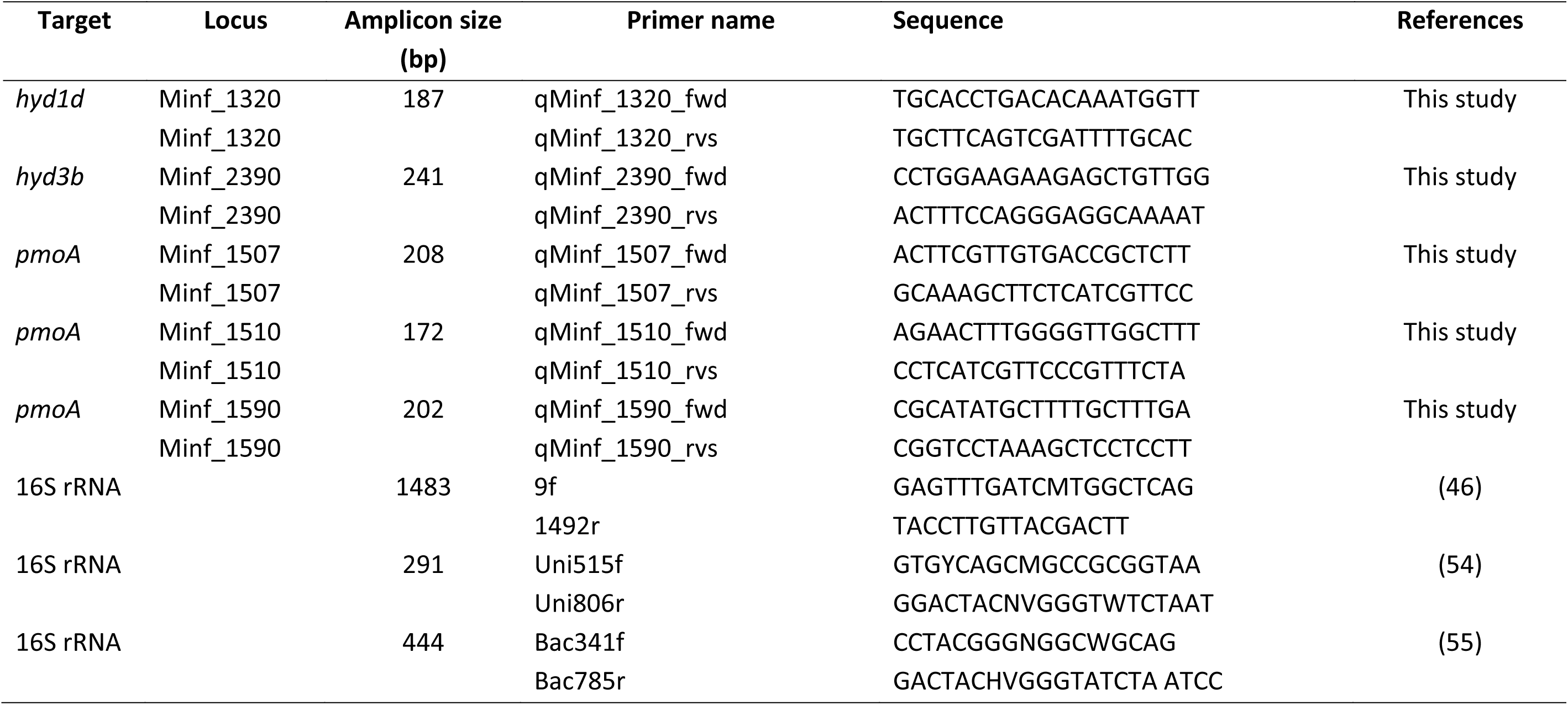
**Table S1.** Primers used for qPCR analysis targeting the methane monooxygenase and hydrogenases of *Methylacidiphilum* spp.

**Table S2:**
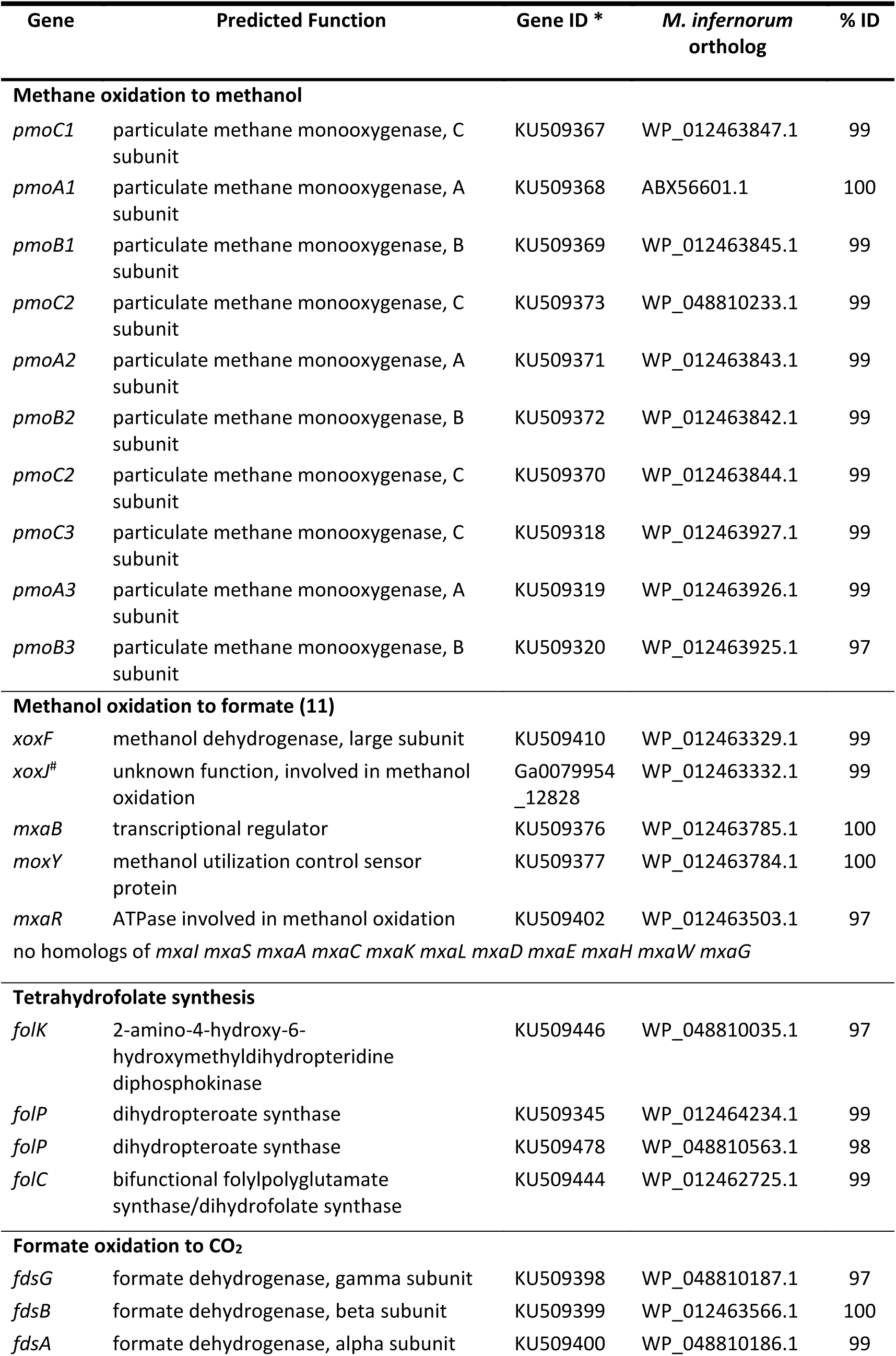
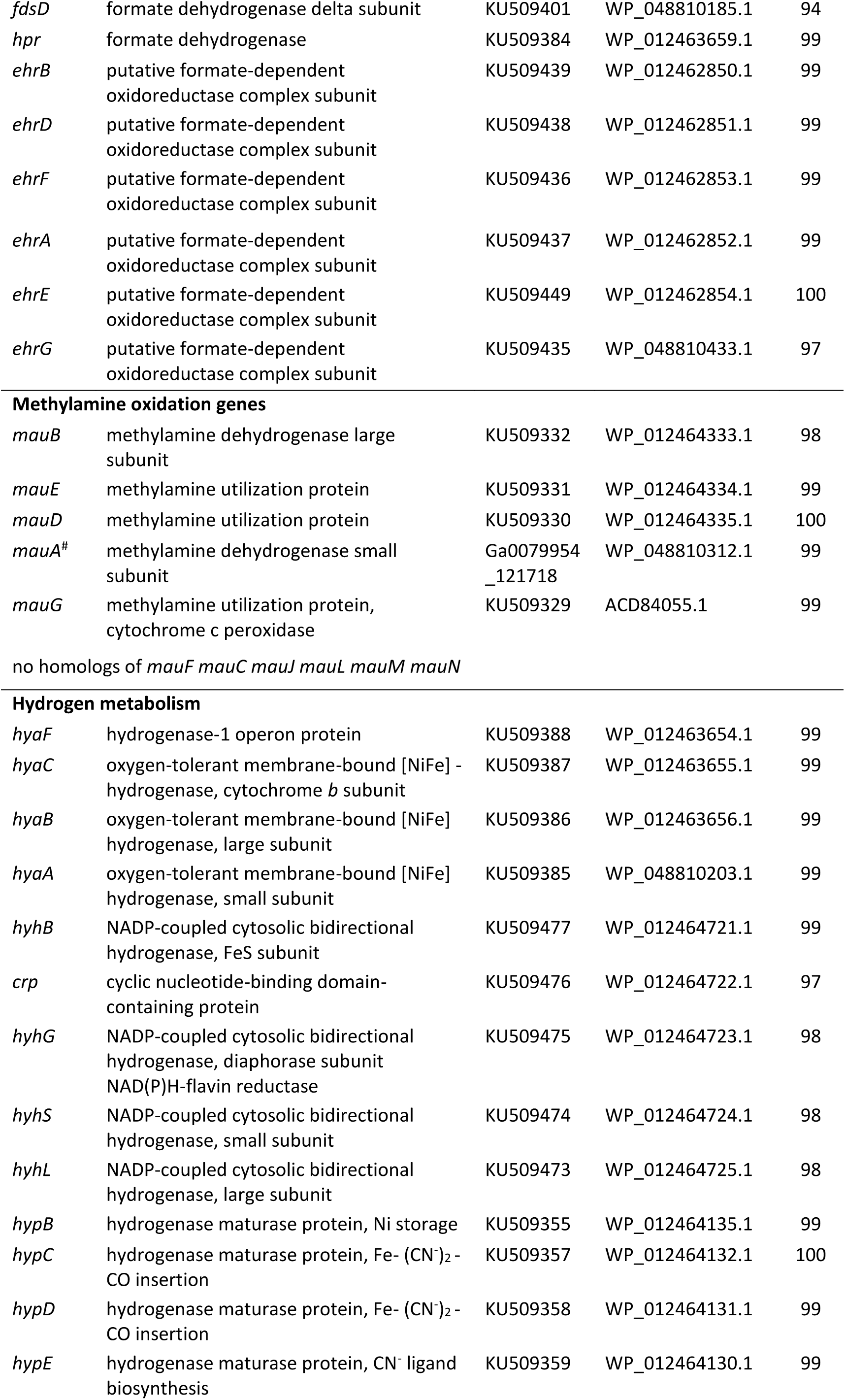
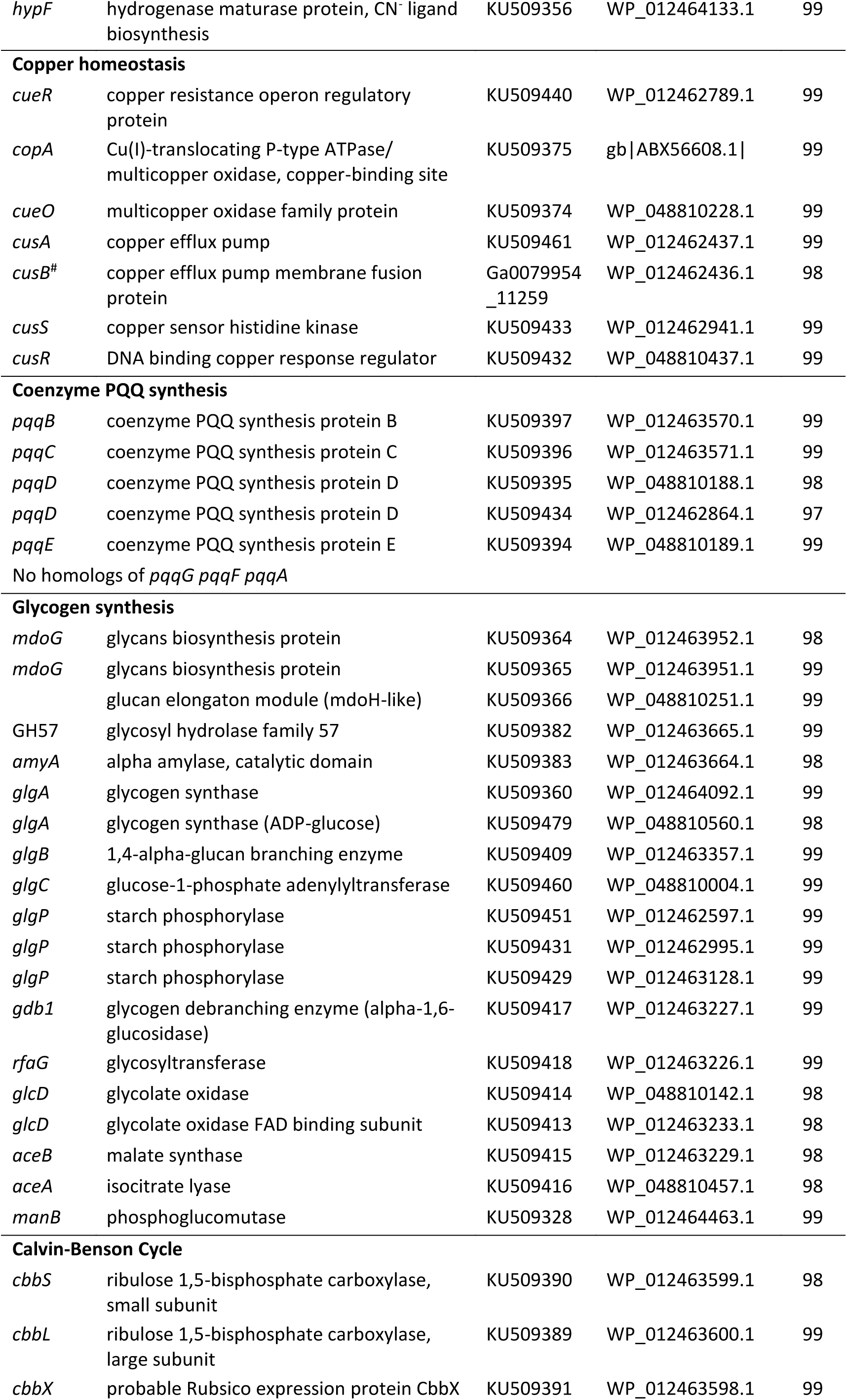
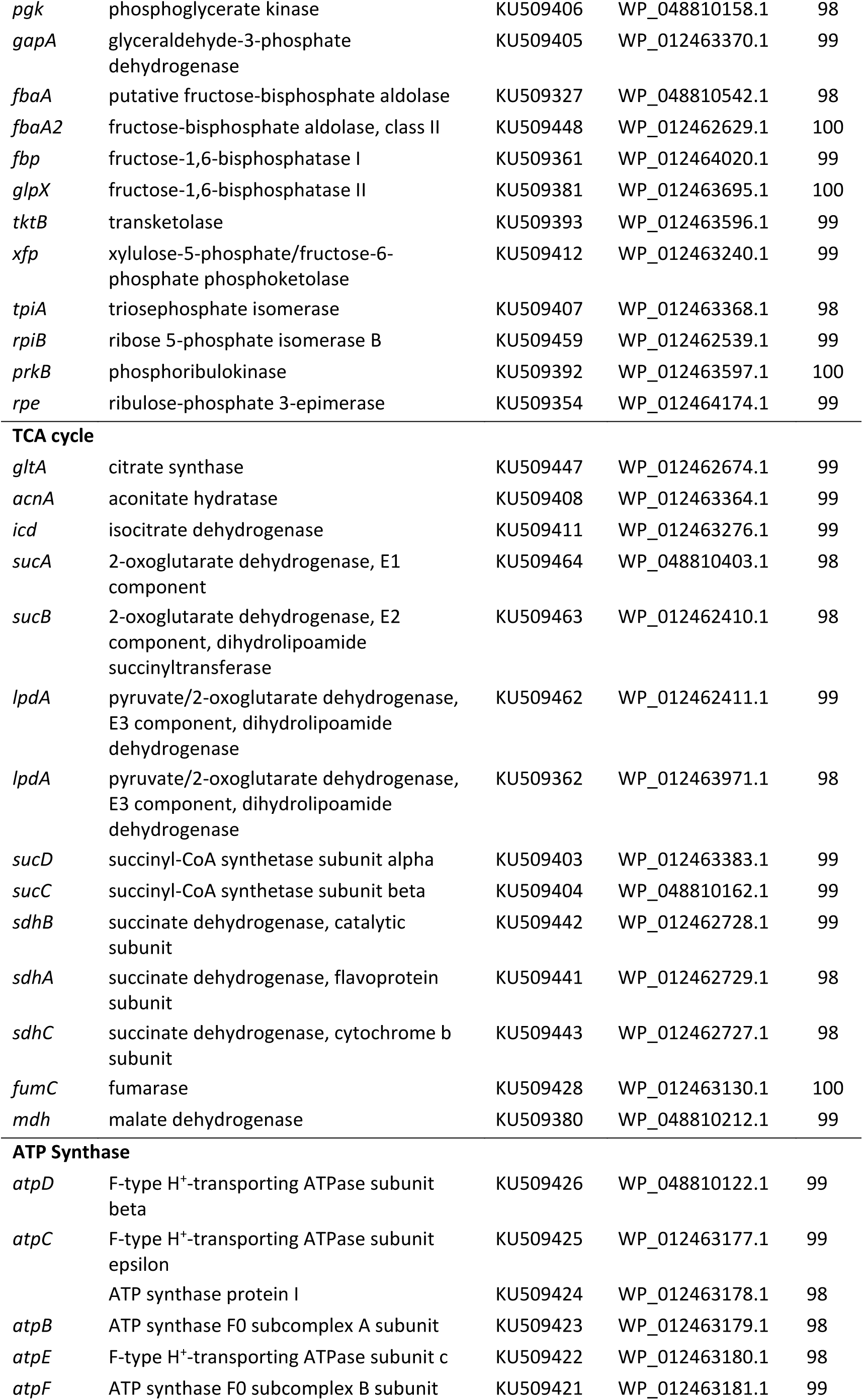
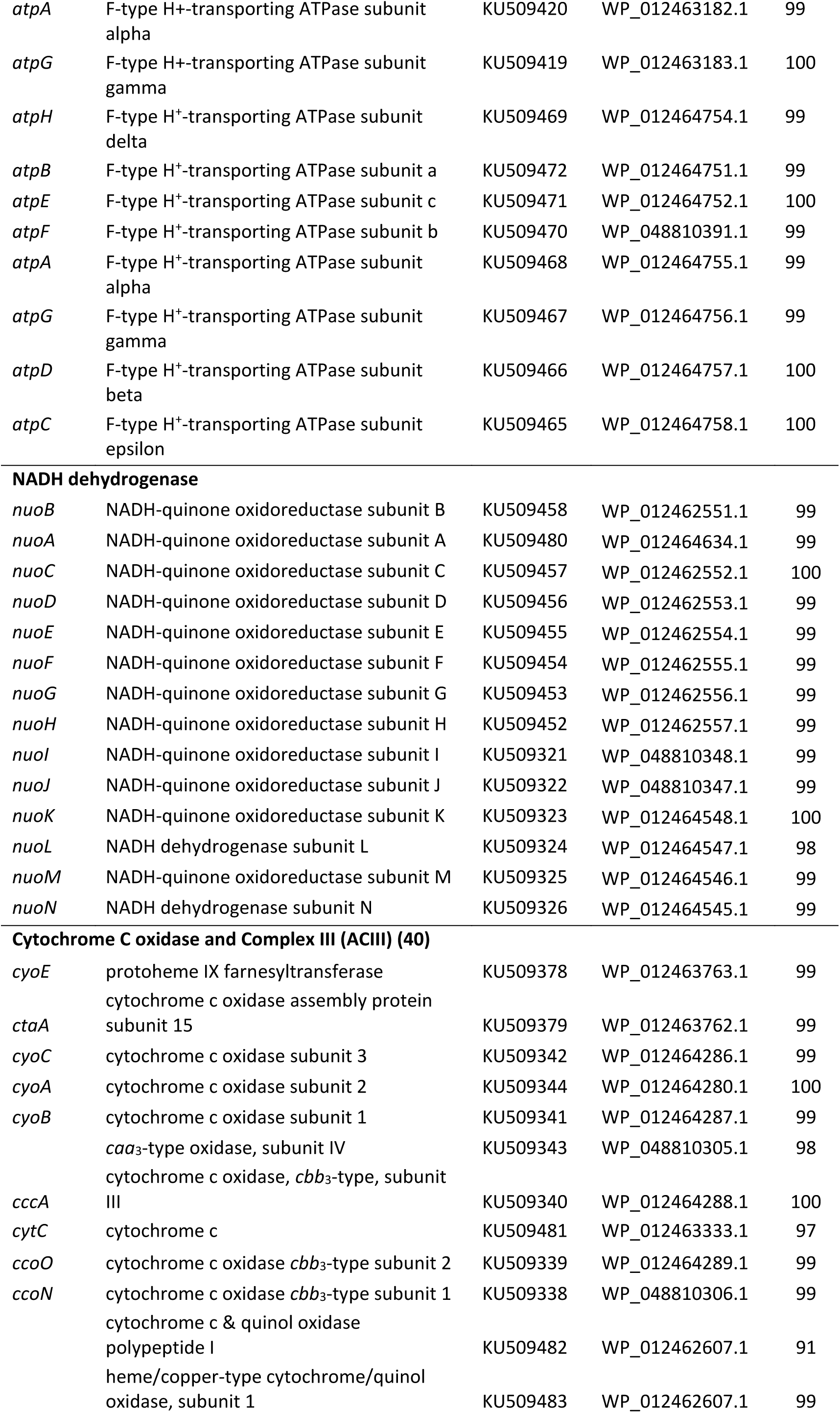
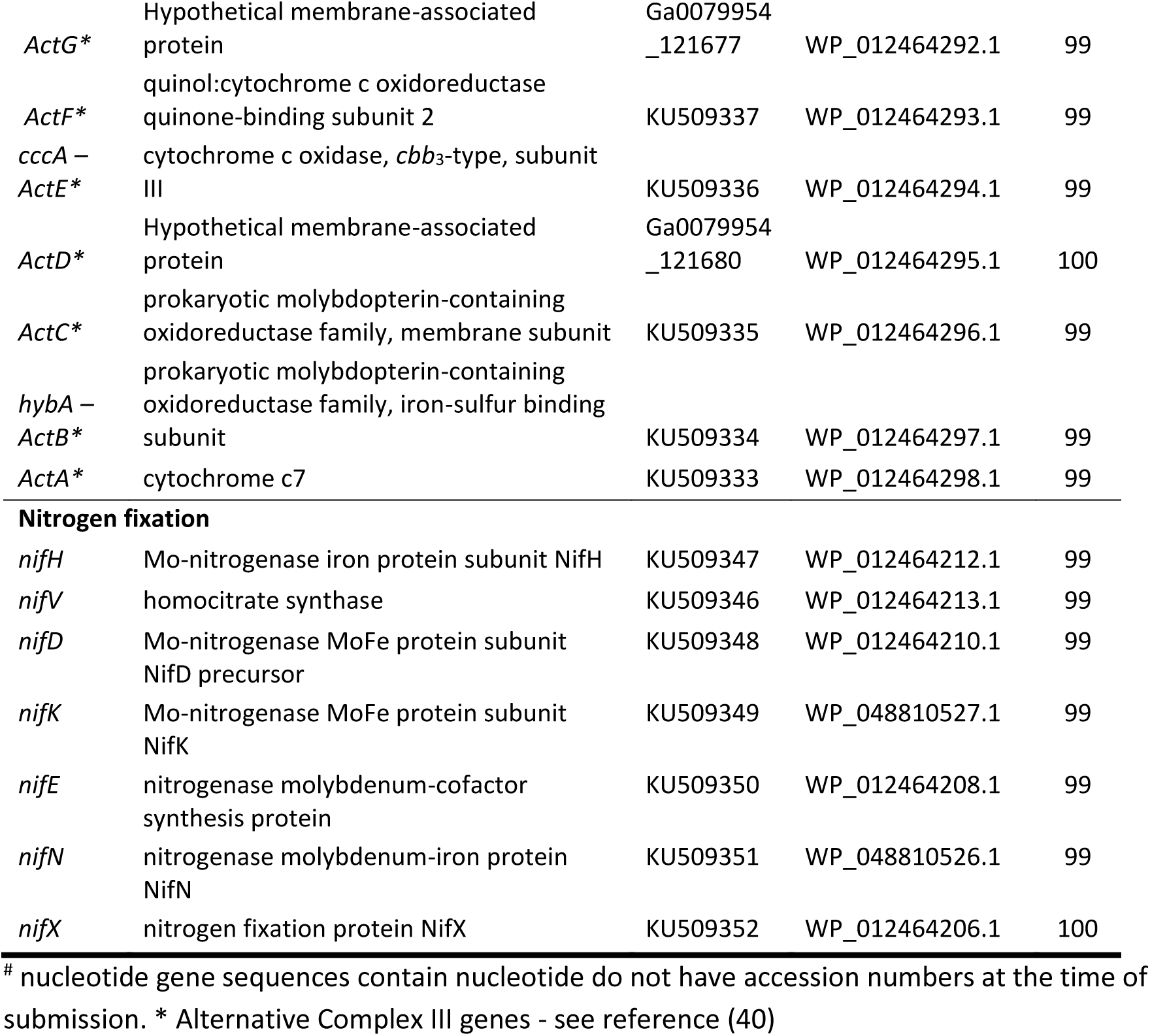
Genes involved in one-carbon and hydrogen metabolism, as well as glycogen storage and utilization in *Methylacidiphilum* sp. RTK17.1. Amino acid orthologs to *Methylacidiphilum infernorum* V4 are shown.

